# Lateral electric field inhibits gel-to-fluid transition in lipid bilayers

**DOI:** 10.1101/2021.01.29.428787

**Authors:** Nidhin Thomas, Ashutosh Agrawal

## Abstract

We report evidence of lateral electric field-induced changes in the phase transition temperatures of lipid bilayers. Our atomic scale molecular dynamics simulations show that lateral electric field increases the melting temperature of DPPC, POPC and POPE bilayers. Remarkably, this shift in melting temperature is only induced by lateral electric field, and not normal electric field. This mechanism could provide new mechanistic insights into lipid-lipid and lipid-protein interactions in the presence of endogenous and exogenous electric fields.

## Introduction

The modulation of physical response of lipid bilayers and cells by electric fields is well documented in the literature. For example, endogenous electric fields have been shown to regulate embryonic development, wound healing, and cancer metastasis [1–4]. Disruption of epithelial layers that form the outer surfaces of skin and organs generates a lateral electric field that may extend over several cell lengths. Epithelial cells sense these electrical signals and respond with directional migration by a process termed electrotaxis [2, 4–7] in order to repair the damaged tissue. Neurites have been shown to grow preferentially towards the negative electrode in the presence of an applied electric field [8]. Electric fields are routinely used to induce electroporation, a phenomenon in which numerous transient pores open up in plasma membrane, furnishing a widely used technique for delivering drugs into cells [9, 10].

Despite the well characterized biological and biomedical roles of electric fields, our understanding of how cells sense and respond to electric fields is still in the formative stages. One identified mechanism is the in-plane migration of charged proteins because of electrophoretic forces. Recently lipid rafts rich in glycolipids have been shown to coalesce and partition membrane proteins in response to electric fields. In this study, we present a new mechanism by which electric field can regulate the physical response of lipid bilayers. Using atomistic studies we show that lateral electric field modulates the gel-to-fluid phase transition temperature of bilayers. This mechanism could have a significant impact on both lipid-lipid and lipid-protein interactions.

## Results

To investigate the effect of lateral electric field, we first simulated a flat bilayer made of 1,2-dipalmitoyl-sn-glycero-3-phosphocholine (DPPC) lipids. DPPC bilayers are the most extensively studied lipid systems, both computationally and experimentally [11, 12]. We performed the simulations in GROMACS 2018 [13] with CHARMM36m [14] force field. The initial structure of the DPPC bilayer was created with 48 lipids per leaflet in CHARMM-GUI[15]. We performed heating and cooling simulations at different electric fields to quantify the changes in the bilayer melting temperature. We obtained the starting structures for heating and cooling simulations by equilibrating bilayers at the starting temperature for at least 200 ns. Heating and cooling scans of equilibrated structures were performed with a rate of ±0.05 K/ns. A slow rate was chosen to ensure minimal impact on the phase transition temperature measurements [12]. The details of the methodology and the simulated systems are presented in the SI.

We use area per lipid (APL) as the core structural measure. The APL is defined as 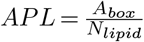, where *A*_*box*_ is the average area of simulation box and *N*_*lipid*_ is the number of lipids per leaflet. Fig. 1 shows the changes in the APL with temperature while heating (red curves) and cooling (blue curves) a bilayer. The changes in the APL are represented by faint background lines but we fit a curve (solid line) to compute the effective change in the APL [12, 16]. The predicted average APL of DPPC lipids is 0.53 nm^2^ at 329 K in the absence of the electric field. This value agrees with an earlier estimate [12], and suggests that the bilayer is in the gel phase. When we increase the temperature to 329.6 K, the APL increases to 0.62 nm^2^. This increase in the APL implies that the bilayer has become fluid and undergone phase transition. The discrete jump observed in the APL curve indicates that the bilayer underwent a first-order phase transition.

**Fig. 1.**
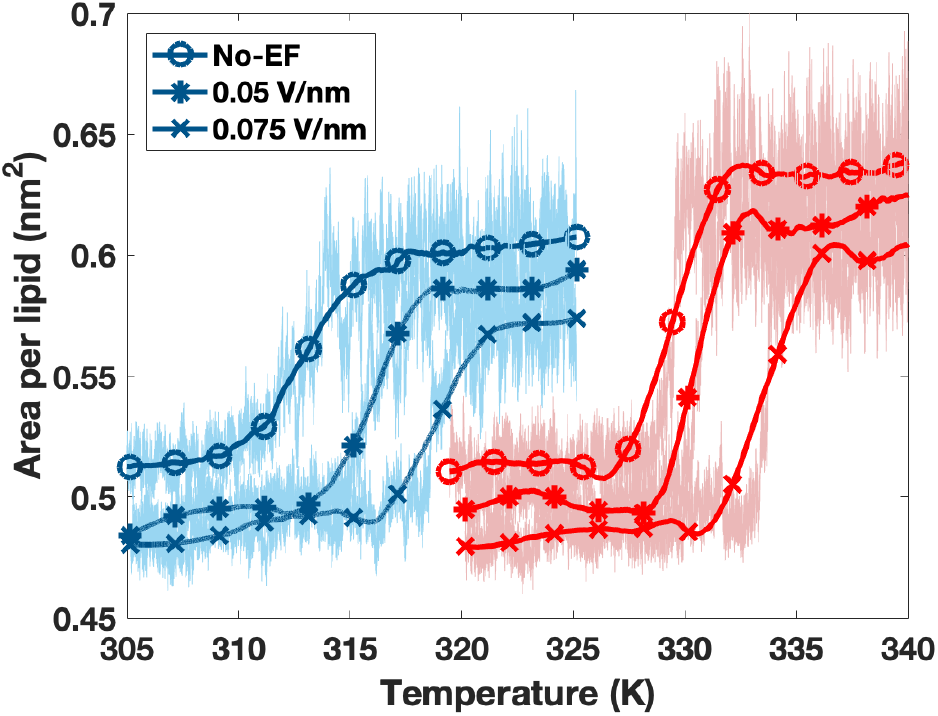
Variation of area per lipid (APL) of DPPC bilayers in heating (red curves) and cooling (blue curves) simulations. The discrete jumps in the red curves depict a first-order phase transition from gel phase to fluid phase. Presence of lateral electric field shifts the discrete jumps rightward, delaying the gel-to-fluid transition. The cooling curves show similar rightward shifts in phase transition temperatures. The relative shifts in the heating and cooling curves show hysteresis emerging from longer time scale associated with disordered state to ordered state transition.

When we subject the bilayer to an electric field of 0.05 V/nm, the APL increases from 0.53 nm^2^ to 0.61 nm^2^ at 330.5 K. When we apply a higher electric field of 0.075 V/nm, the APL jumps from 0.53 nm^2^ to 0.60 nm^2^ at 333.9 K. Thus, the lateral electric fields shift the melting temperature by 1.0 K and 4.5 K for the DPPC bilayer. Fig. 2a shows the DPPC bilayer at 333 K in the absence of the electric field. The bilayer is in the fluid phase and has a thickness of 4.0 nm. Fig. 2b shows the bilayer at the same temperature but in the presence of 0.075 V/nm lateral electric field. The lipid acyl chains become straighter and the bilayer thickness increases to 4.4 nm as the bilayer is in the gel phase because of delayed melting induced by the electric field.

**Fig. 2.**
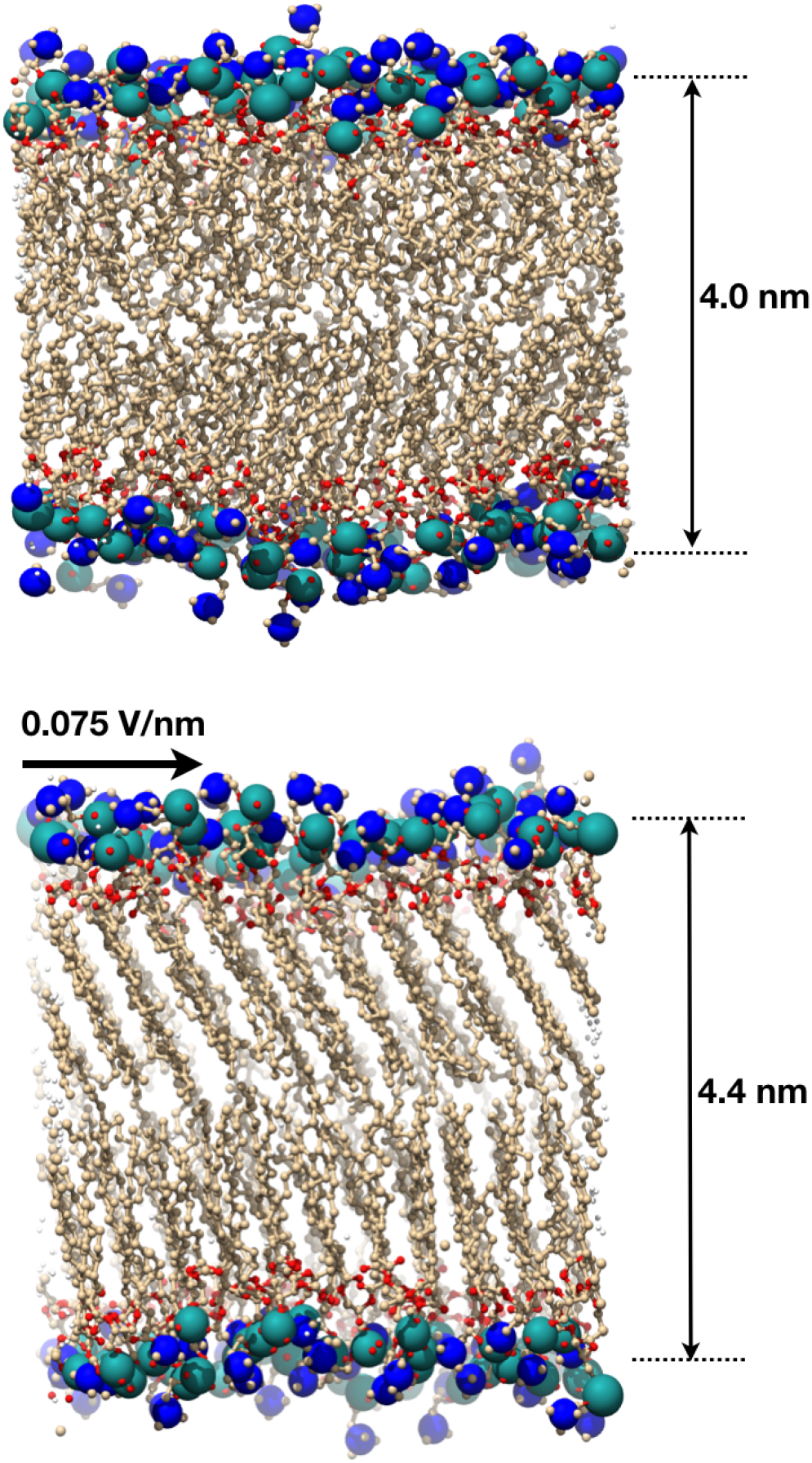
DPPC bilayer at 330 K in the absence and presence of electric field. (a) DPPC bilayer is in the fluid phase in the absence of the applied electric field. The acyl chains are disordered and the bilayer has a thickness of 4 nm. (b) The bilayer is in the gel phase at the same temperature in the presence of a lateral electric field of 0.075 V/nm. The lipids are straighter and the bilayer thickness is increases to 4.4 nm.

The reverse transition curves obtained from cooling simulations are shifted to lower melting temperatures and show hysteresis. Phase transition occurs at 312.6 K, 316.2 K and 318.8 K, respectively for 0 V/nm, 0.05 V/nm and 0.075 V/nm electric fields. The shifted transition temperatures and the hysteresis occur because the process of ordering from a disordered state occurs on a slower time scale compared to the process of disordering from an ordered state. While gel-to-fluid transition occurs via a one-step process, fluid-to-gel transition has been shown to get trapped in ripple-like phases creating hysteresis [12]. Because of this difference in the time scales associated with the heating and the cooling simulations, heating runs provide a more accurate quantitative measure of the phase transition boundaries. Nonetheless, cooling runs provide a qualitative validation of the results.

To verify our findings further, we computed the fraction of the acyl chain gauche dihedrals as the second structural measure. The gauche angle is the fraction of the dihedral angle of acyl chain carbon atoms(*ψ*) which are within −60° and 60°, computed directly from GROMACS. Fig. 3 shows the evolution of the fraction of the gauche angle with temperature. At lower temperatures, the DPPC membrane has a small fraction of gauche angle indicating the gel phase with acyl chains in the *trans* configuration. As the temperature increases, the ordered lipid acyl chains start to melt and undergo gauche rotational isomerization. As a result, the fraction undergoes a jump at the phase transition temperature confirming the predictions revealed by the APL plots. The heating plots show phase transition at 329.5 K and 333.5 K for 0 V/nm and 0.075 V/nm. A jump in the gauche angle can also be observed in the cooling simulations. Fig. 3 also shows the hysteresis behaviour during the heating and the cooling simulations confirming the first-order phase transition in DPPC bilayers.

**Fig. 3.**
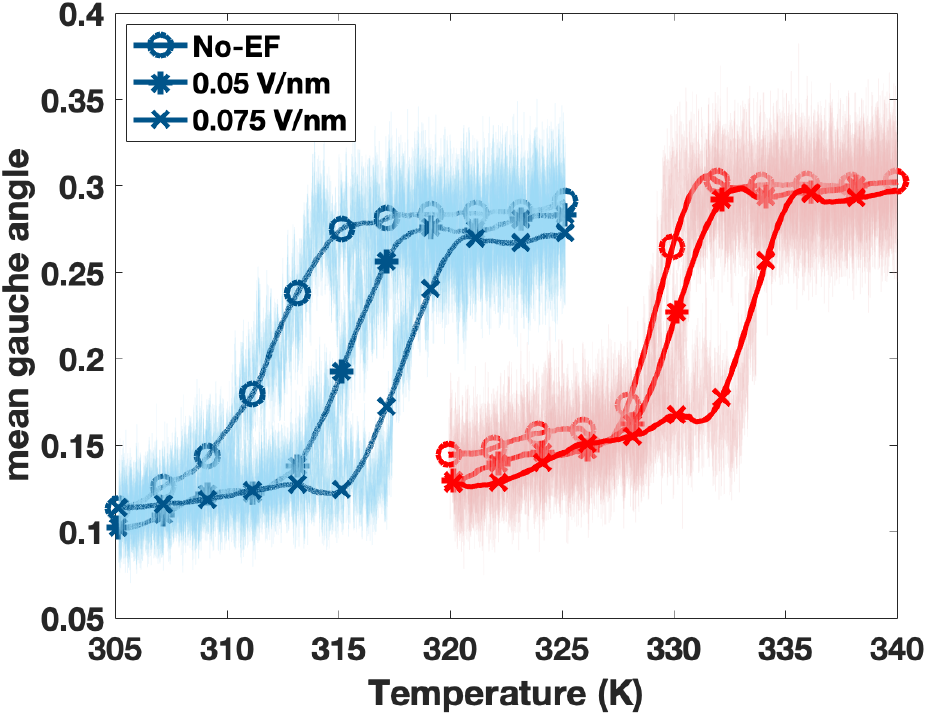
Fraction of gauche angle of acyl chains in the DPPC bilayer. The heating (red curves) and cooling (blue curves) calculations confirm the electric-field induced shifts in the phase transition temperature.

Having confirmed the effect of lateral electric on DPPC bilayers, we simulated its effect on two other key lipids: 1-palmitoyl-2-oleoyl-sn-glycero-3-phosphocholine (POPC) and 1-palmitoyl-2-oleoyl-sn-glycero-3-phosphoethanolamine (POPE). Fig. 4 shows the APL plots as a function of temperature and electric field. The APL increases from 0.54 nm^2^ to 0.63 nm^2^ at 275 K temperature for the POPC bilayer in the absence of the electric field. After subjecting the bilayer to an electric field of 0.075 V/nm, the APL increased from 0.52 nm^2^ to 0.60 nm^2^ at 288 K. Thus, the lateral electric field induces a 13 K shift in the phase transition temperature of the POPC bilayer. Similarly, for POPE bilayer, the APL increases from 0.49 nm^2^ to 0.59 nm^2^ at 316 K in the absence of the electric field. In the presence of 0.075 V/nm electric field, the APL increases from 0.48 nm^2^ to 0.58 nm^2^ at 332 K. Thus, the lateral electric field shifts melting temperature by 16 K for the POPE bilayer. These results demonstrate that lateral electric field affects a wide variety of lipids. They also show that the effect of lateral electric field is different for different lipid types. The shifts obtained for the POPC and POPE bilayers are much larger than that obtained for DPPC bilayer. These predictions for the three lipid systems are confirmed by repeated simulations with different starting structures presented in Figs. S1 and S2.

**Fig. 4.**
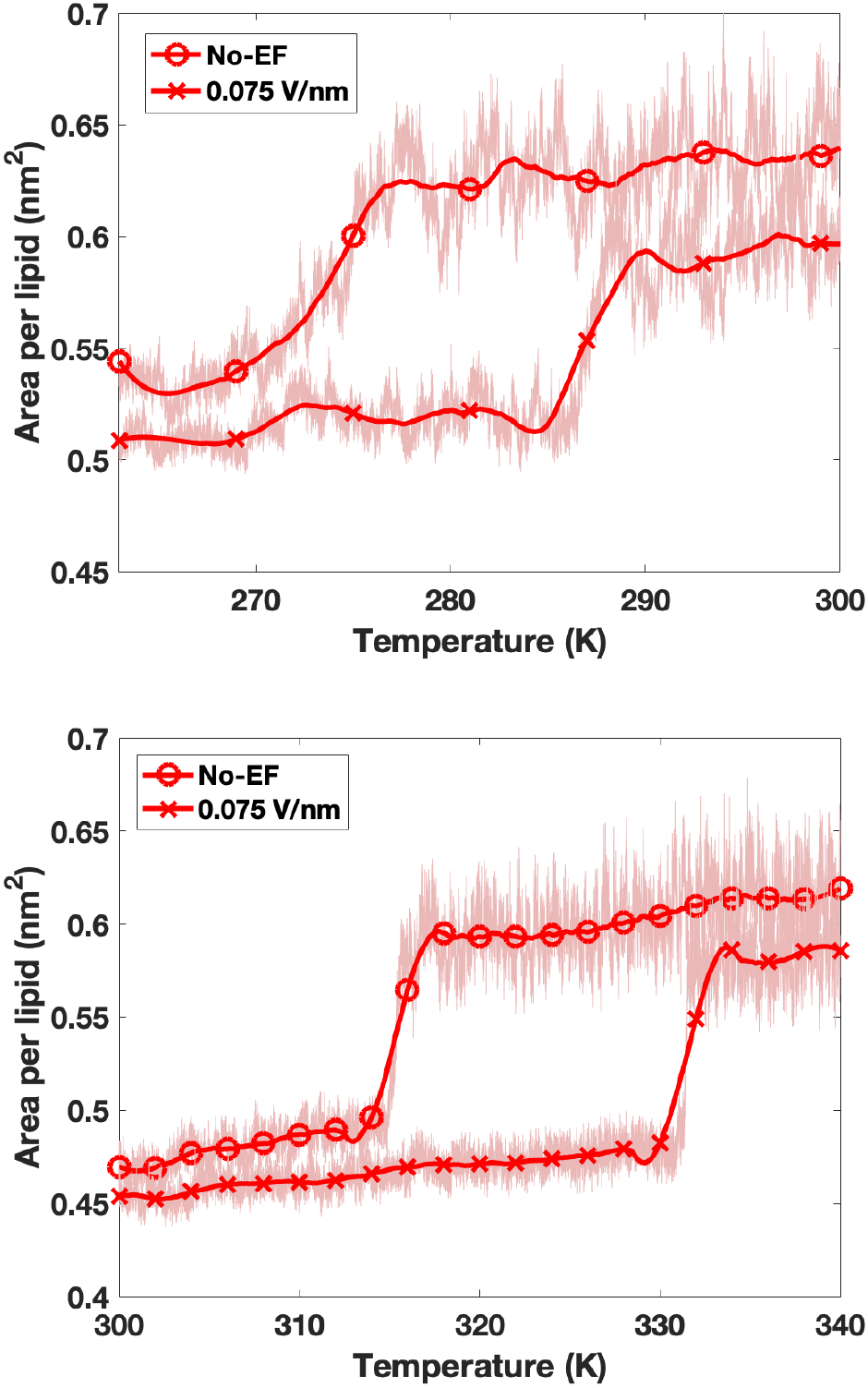
Variation of area per lipid (APL) of POPC (a) and POPE (b) bilayers during heating simulations. The curves in the presence of electric field show significant rightward shifts in the phase transition temperatures.

## Discussion

Endogenous and exogenous electric fields play a critical role in growth and repair of cells and tissues. In this study, we present a new mechanism by which electric fields can control the physical response of cellular membranes. Our findings show that lateral electric fields can regulate the phase transition in lipid bilayers in lipid-dependent manner. As cellular membranes invariably have heterogeneous lipid composition and electric fields possess spatial variations, cellular membranes can have a heterogeneous distribution of gel and fluid phases. This electric-field dependent property can contribute to the directional response of cells in the presence of electric fields. These local phase changes can also have a significant impact on membrane-protein interactions.

We can understand the electric field-induced phase changes by its impact on the bilayer structure. Lateral electric field reorients the P-N dipoles in the lipid headgroups. This generates an in-plane polarization in the bilayer. According to a recent electromechanical theory [17], the in-plane polarization generated by a tangential electric field generates a compressive stress in a bilayer. This compressive stress in turn would shift the gel-to-fluid phase transition to a higher temperature. The inhibition of gel-to-fluid transition by compressive stress has been demonstrated in previous atomistic studies [18, 19]. The presence of compressive stress in the bilayer is also confirmed by Fig. 5 which shows the APL of the three lipids in the fluid phase. The histograms in the presence of electric field are shorter than the those in the absence of electric field. Thus, despite being in the same phase, the lipids have smaller APLs in the presence of an electric field indicating the existence of compressive stress in the bilayers.

**Fig. 5.**
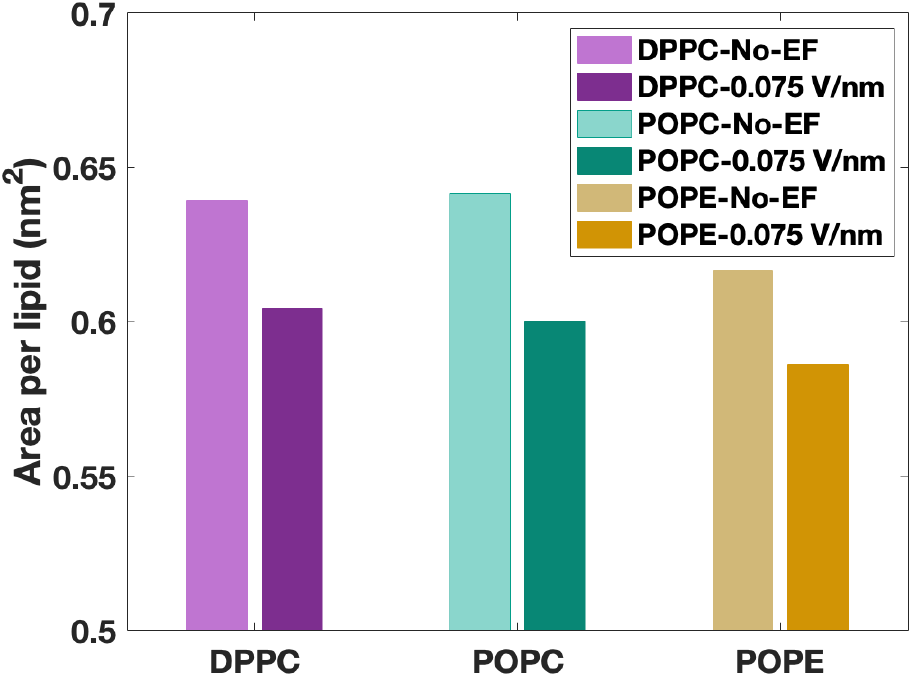
Lateral electric field reduces the APL of the bilayer systems in the fluid phase. Despite being in the same phase, bilayers with and without electric field have different APLs. This change is a consequence of in-plane compression generated in the bilayer by the lateral electric field.

Finally, we would like to emphasize that the phase change presented in this study is only induced by lateral electric field. Application of normal electric field does not lead to any noticeable shift in the phase transition temperatures. Fig. 6 shows the heating and cooling simulations for DPPC lipids in the presence of 0 V/nm and 0.075 V/nm electric field perpendicular to the bilayer. The two sets of plots almost overlap showing no shifts in the meting temperature. While a majority of the theoretical and computational investigation has been performed with normal electric fields, our work with lateral electric field and its consequences offers a fresh perspective into the electromechanical behaviour of cellular membranes.

**Fig. 6.**
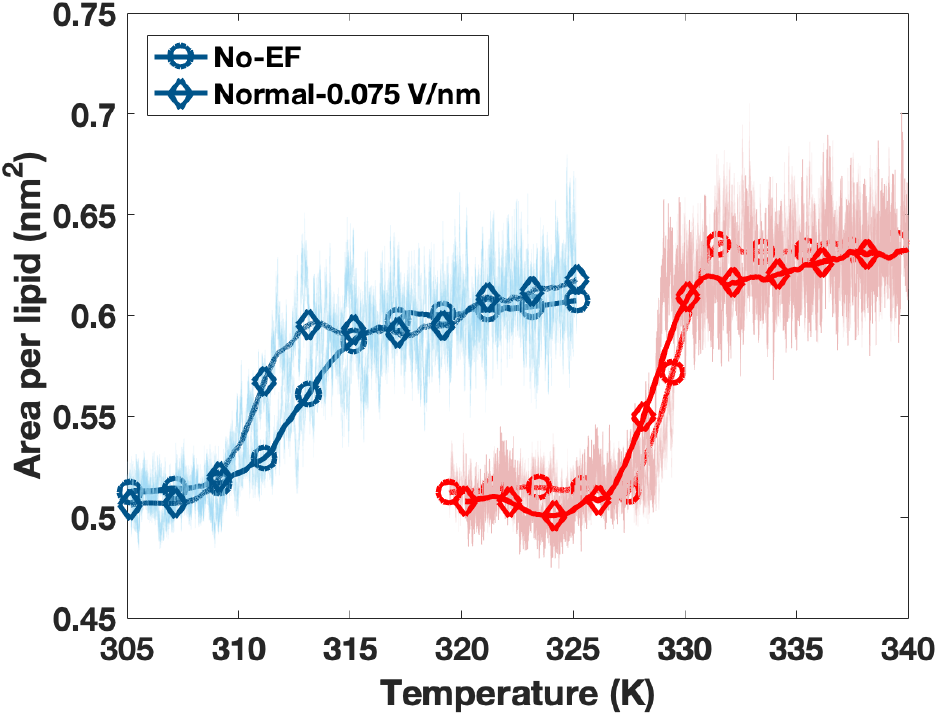
Normal electric field does not shift phase transition temperature in DPPC bilayer. The phase transition shift is uniquely induced by lateral electric field.

## ACKNOWLEDGEMENTS

A.A. acknowledges support from NSF Grants CMMI-1727271 and CMMI-1931084. The authors acknowledge the use of the Opuntia, Sabine, and Carya clusters to perform the simulations and the advanced technical support from the Research Computing Data Core at UH to carry out the research presented here.

