## Supplemental Document for "Lateral electric field inhibits gel-to-fluid transition in lipid bilayers"

### Supporting Information for "Lateral electric field inhibits gel-to-fluid transition in lipid bilayers"

**[1] Department of Mechanical Engineering, University of Houston, Houston, TX 77204**

**Corresponding authors:**
*****

### Molecular dynamics protocol

All-atom models of DPPC, POPC and POPE lipids were chosen to run molecular dynamics simulations. In the first phase, equilibrium simulations of DPPC lipids were performed at different lateral electric field intensities and temperatures. Lipid bilayer structural properties of DPPC lipids obtained from these equilibrium simulations were analyzed. These analyses showed that lateral electric field modifies the structure of lipid bilayers and alter the phase transition temperature of DPPC bilayer. In order to ascertain this hypothesis, we performed heating and cooling simulations such that we could precisely measure the phase transition temperatures at various lateral electric field intensities. In addition, we performed heating and cooling simulations on POPC and POPE lipids to verify the shift in phase transition temperatures and understand how lipid acyl tails and headgroup dipoles impact in the phase transition temperatures. Simulated annealing simulations were performed with zero external electric field and 0.075 V/nm external electric field. These simulations were performed at 0.05 K/ns rate. Details of the simulations are provided in Table 1. Each simulation was reproduced to verify the results.

#### Equilibrium simulations

Initial structure of DPPC all-atom model was created using CHARMM-GUI [1]. Bilayer system consisted of 48 lipids per leaflet and 50 water molecules per lipid. Salt ions were not added as DPPC lipids are neutral and lateral electric field may produce concentration gradient across the bilayer. Simulations were performed in GROMACS 2018 [2] using CHARMM36m [3] force field. We performed energy minimization using steepest descent algorithm with maximum force limit of 700 KJ/mol.nm. It was followed by three steps of NVT equilibration and three steps of NPT equilibration covering a total of 500 ps before NPT production runs. Throughout the simulation, system was maintained at fixed temperature and pressure at 1 atm. Details of the temperature of each system and total production run time is mentioned in Table 2. During equilibration, Z-position of phosphorus atoms of lipids were constrained in order to maintain the bilayer structure till the system settle down. These constraints were gradually removed in each step of equilibration. Berendsen thermocouple [4] with 1.0 ps time constant and Berendsen semi-isotropic pressure couple with 5.0 ps time constant and 4.5 $\times 10^5$ bar $^{-1}$ compressibility were used throughout the equilibration. In production run, Nosé-Hoover thermocouple [5] and Parrinello-Rahman pressure couple [6] were used. Constant temperature equilibrium simulations at below the phase transition temperatures were performed with initial structure obtained from cooling simulations. However, those structures were also equilibrated prior to production runs. We used Verlet cut-off scheme used throughout the simulation. Van der Waals interactions were cut-off at 1.2 nm and used force-switch vdw-modifier at 1.0 nm. Coulombic interactions were cut-off at 1.2 nm and reciprocal space interactions were calculated using PME [7]. LINCS algorithm [8] was used to constrain hydrogen bonds.External electric fields were applied in Y-direction, parallel to the bilayer surface. Production runs were performed for at least 300 ns and only last 200 ns simulation data was used to analyze the data and earlier frames were discarded as part of equilibration.

#### Area per lipid calculations

Area per lipid (APL) was computed by following formula.

$$APL = \frac{L_x.L_y}{N_l} \qquad [S1]$$

where $L_x$ and $L_y$ are the dimension of simulation box in X and Y direction and $N_l$ is the number of lipids per leaflet.

#### Gauche angle calculations

Fractional gauche angles is another method to quantify the ordering effect of lipid acyl tails. Dihedral angle ($\phi$) '1' corresponds to C21-C22-C23-C24 carbon atoms of lipid tails. Similarly, C213-C214-C215-C216 angle of acyl tails correspond to dihedral angle '13'. 'gmx angle' command of GROMACS was used to compute the fraction of gauche angles. Gauche angle is defined as the $-60 < \phi < 60$. For equilibrium simulations, fractional gauche angle of lipid acyl tails were measured for bilayer systems at 323K and varying external electric field intensities. Comparison of fraction of gauche angles show clearly that application of external electric field reduce the fraction of gauche angle and thereby increase the order of tails. In addition, applied electric field intensity of 0.10 V/nm lower the gauche angle fraction significantly denoting the phase transition effect. Therefore, it can be deduced that DPPC bilayer at 323K remain to be at $l_o$ phase when 0.10 V/nm is applied in comparison to DPPC bilayer at no external electric field. Phase transition of lipid bilayers in simulated annealing systems were measured by averaging individual dihedral angles per frame. The difference in phase transition points in heating and cooling simulations suggests the hysteresis behaviour of bilayer systems.

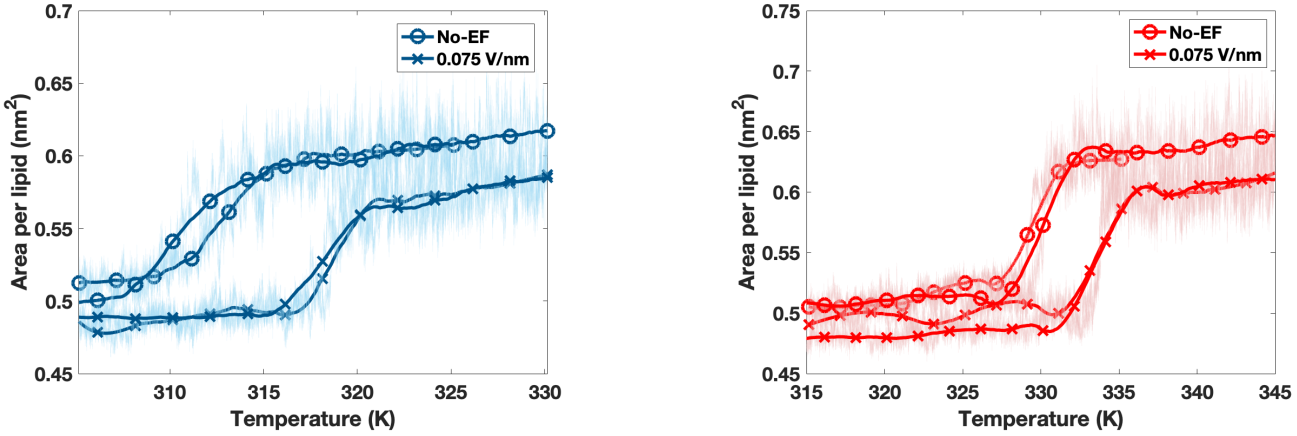


Figure S1: APL evolution obtained from repeat (**a**) cooling and (**b**) heating simulations of DPPC lipid bilayer. Details of the phase transition temperatures are provided in Table 3. Phase transition temperature was shifted by ∼6 K and ∼4.5 K in cooling and heating simulations, respectively upon application of 0.075 V/nm external electric field.

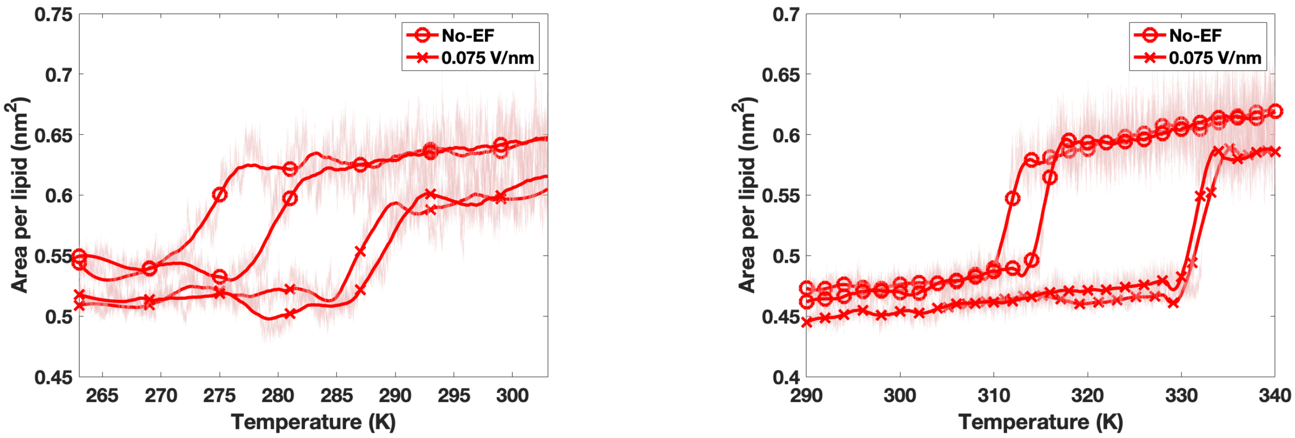


Figure S2: APL evolution obtained from repeat heating simulations of (**a**) POPC and (**b**) POPE lipid bilayers. Details of the phase transition temperatures are provided in Table 3. Phase transition temperature was shifted by ∼13 K and ∼16 K in POPC and POPE simulations, respectively upon application of 0.075 V/nm external electric field.

| System | Lipids per leaflet | Rate (K/ns) | Time (ns) | EF (V/nm) | Number of simulations |
|---|---|---|---|---|---|
| DPPC | 48 | +0.05 | 800 | 0.0 | 2 |
| DPPC | 48 | +0.05 | 1000 | 0.075 | 2 |
| DPPC | 48 | -0.05 | 800 | 0.0 | 2 |
| DPPC | 48 | -0.05 | 1000 | 0.075 | 2 |
| POPC | 48 | +0.05 | 1000 | 0.0 | 2 |
| POPC | 48 | +0.05 | 1000 | 0.075 | 2 |
| POPE | 48 | +0.05 | 1000 | 0.0 | 2 |
| POPE | 48 | +0.05 | 1000 | 0.075 | 2 |
| DPPE | 48 | +0.05 | 1000 | 0.0 | 2 |
| DPPE | 48 | +0.05 | 1000 | 0.075 | 2 |
| DPPC-Normal EF | 48 | +0.05 | 1000 | 0.075 | 2 |
| DPPC-Normal EF | 48 | -0.05 | 1000 | 0.075 | 2 |

Table 1: Summary of heating (+ sign) and cooling (− sign) simulations. Electric field was applied laterally, parallel to the bilayer-water interface, except for Normal-EF systems in which electric field was applied perpendicular to the bilayer-water interface.

| System | Lipids per leaflet | Time (ns) | Temperature (K) | EF (V/nm) |
|---|---|---|---|---|
| DPPC | 48 | 300 | 293 | 0.0 |
| DPPC | 48 | 300 | 303 | 0.0 |
| DPPC | 48 | 300 | 313 | 0.0 |
| DPPC | 48 | 300 | 323 | 0.0 |
| DPPC | 48 | 300 | 333 | 0.0 |
| DPPC | 48 | 300 | 343 | 0.0 |
| DPPC | 48 | 300 | 293 | 0.05 |
| DPPC | 48 | 300 | 303 | 0.05 |
| DPPC | 48 | 300 | 313 | 0.05 |
| DPPC | 48 | 300 | 323 | 0.05 |
| DPPC | 48 | 300 | 333 | 0.05 |
| DPPC | 48 | 300 | 343 | 0.05 |
| DPPC | 48 | 300 | 293 | 0.075 |
| DPPC | 48 | 300 | 303 | 0.075 |
| DPPC | 48 | 300 | 313 | 0.075 |
| DPPC | 48 | 300 | 323 | 0.075 |
| DPPC | 48 | 300 | 333 | 0.075 |
| DPPC | 48 | 300 | 343 | 0.075 |
| DPPC | 48 | 300 | 293 | 0.10 |
| DPPC | 48 | 300 | 303 | 0.10 |
| DPPC | 48 | 300 | 313 | 0.10 |
| DPPC | 48 | 300 | 323 | 0.10 |
| DPPC | 48 | 300 | 333 | 0.10 |
| DPPC | 48 | 300 | 343 | 0.10 |
| POPC | 48 | 400 | 253 | 0.0 |
| POPC | 48 | 400 | 253 | 0.075 |
| POPE | 48 | 400 | 290 | 0.0 |
| POPE | 48 | 400 | 290 | 0.075 |
| DPPC-Normal EF | 48 | 400 | 345 | 0.0 |
| DPPC-Normal EF | 48 | 400 | 345 | 0.075 |

Table 2: Summary of equilibrium simulations at constant temperature. Electric field was applied laterally, parallel to the bilayer-water interface, except for Normal-EF systems in which electric field was applied perpendicular to the bilayer-water interface. More lipids were used in normal EF systems since smaller patches do not show electroporation associated with normal-EF.
